# Genome-wide measurement of spatial expression in patterning mutants of *Drosophila melanogaster*

**DOI:** 10.1101/046128

**Authors:** Peter A. Combs, Michael B. Eisen

## Abstract

Genome sequencing has become commonplace, but the understanding of how those genomes ultimately specify cell fate during development is still elusive. Extrapolating insights from deep investigation of a handful of developmentally important Drosophila genes to understanding the regulation of all genes is a major challenge. The developing embryo provides a unique opportunity to study the role of gene expression in pattern specification; the precise and consistent spatial positioning of key transcription factors essentially provides separate transcriptional-readout experiments at a critical point in development.

We cryosectioned and sequenced mRNA from single Drosophila melanogaster embryos at the blastoderm stage to screen for spatially-varying regulation of transcription. Expanding on our previous screening of wild type embryos, here we present data from dosage mutants for key maternally provided regulators, including depletion of zelda and hunchback and both over-expression and depletion of bicoid. These data recapitulate all of the expected patterning changes driven by these regulators; for instance, we show spatially-confined up-regulation of expression in the bicoid over-expression condition, and down-regulation of those genes in the bicoid knock-down case, consistent with bicoid’s known function as an anterior-localized activator.

Our data highlight the role of combinatorial regulation of patterning gene expression. When comparing changes in multiple conditions, genes responsive to one mutation tend to respond to other mutations in a similar fashion. Furthermore, genes that respond differently to these mutations tend to have more complex patterns of TF binding.

## Introduction

Proper animal development relies on complex, highly coordinated gene expression patterns in both space and time. In *Drosophila* (and many other long germ-band insects), this is achieved through nuclei nearly simultaneously reading out a number transcription factors (TFs) that have been maternally deposited in a spatially dependent manner, which establishes the dorsal-ventral and anterior-posterior axes. In many eukaryotes, this readout of regulatory signals is mediated through enhancers and other cis-regulatory elements that are located anywhere from several kilobases to megabases away from the promoters of the genes they regulate.

The classical reverse genetics approach involves mutating a gene, then investigating the phenotypic consequences of that mutation. This approach has been extremely successful in helping to elucidate the mechanisms of transcriptional regulation for a number of key loci. However, historically *in situ* hybridization has been used to assay these consequences. By necessity, previous studies have focused only on the patterning changes of a handful of genes, since assaying more genes is impractical. While these changes are often rationally selected—including known transcription factors that, in turn, drive other patterns—the scale of previous experiments means that they have only illustrated the common types of changes observed, rather than completely cataloging a given TF’s effects.

For the TFs we have investigated in this study (*bicoid*, *hunchback,* and *zelda),* there are high quality chromatin immunoprecipitation (ChIP) experiments already in the literature for wild-type flies (MacArthur et al. 2009; Harrison et al. 2011). These experiments can suggest potential targets, but due to the large number of binding sites in the genome as well as the dense, interconnected regulatory network that can potentially buffer changes, simply performing *insitus* on every gene with a binding site is a prohibitively large experiment, and unlikely to perform radically better than chance. Strong binding is suggestive, but not conclusive, for functional effect. TFs bind to hundreds of sites throughout the genome, and can show strong ChIP signal at many hundreds more where no binding is found. The true gold-standard for assaying function of a particular binding site must therefore demonstrate some downstream effect that is different when that binding is removed.

In this study, we extend our previous method for sequencing mRNA from spatially restricted *Drosophila melanogaster* samples and apply it to a number lines that are mutant for TFs known to be important in establishing proper spatial patterns of expression. Our goal was to determine which patterned genes show distinct, TF dependent spatio-temporal regulation at the blastoderm stage. We identify a large number of genes with both expected and unexpected patterning changes, and through integrated analysis of previous ChIP data for a large number of anterior-posterior patterning factors, we highlight common themes in genes with complex dependence on these TFs.

## Results

### Genome-wide atlases of the blastoderm stage of multiple dosage mutants

We sliced embryos and sequenced the resulting mRNA from 4 mutant genotypes (fig. 1A): a *zld* germline clone, an RNAi knockdown for *bcd,* a knockdown for *hb,* and an overexpression line for *bcd* with approximately 2.4× wildtype expression. We chose two time points: cycle 13 (determined using nuclear density of either DAPI stained embryos or of the Histone-RFP present in the *zld* line) and mid-to-late cycle 14 (determined using 50-65% membrane invagination at stage 5) (fig. 1B). Genes expressed in cycle 13 are towards the end of the early round of genome activation and are enriched for *ZLD* binding (Tadros and Lipshitz 2009; Lott et al. 2011; Harrison et al. 2011), but are early enough that the majority of patterning disruptions are likely to be direct effects of the mutants. By contrast, we chose the stage 5 time point in order to highlight the full extent of the patterning changes across the network.

**Figure 1.**
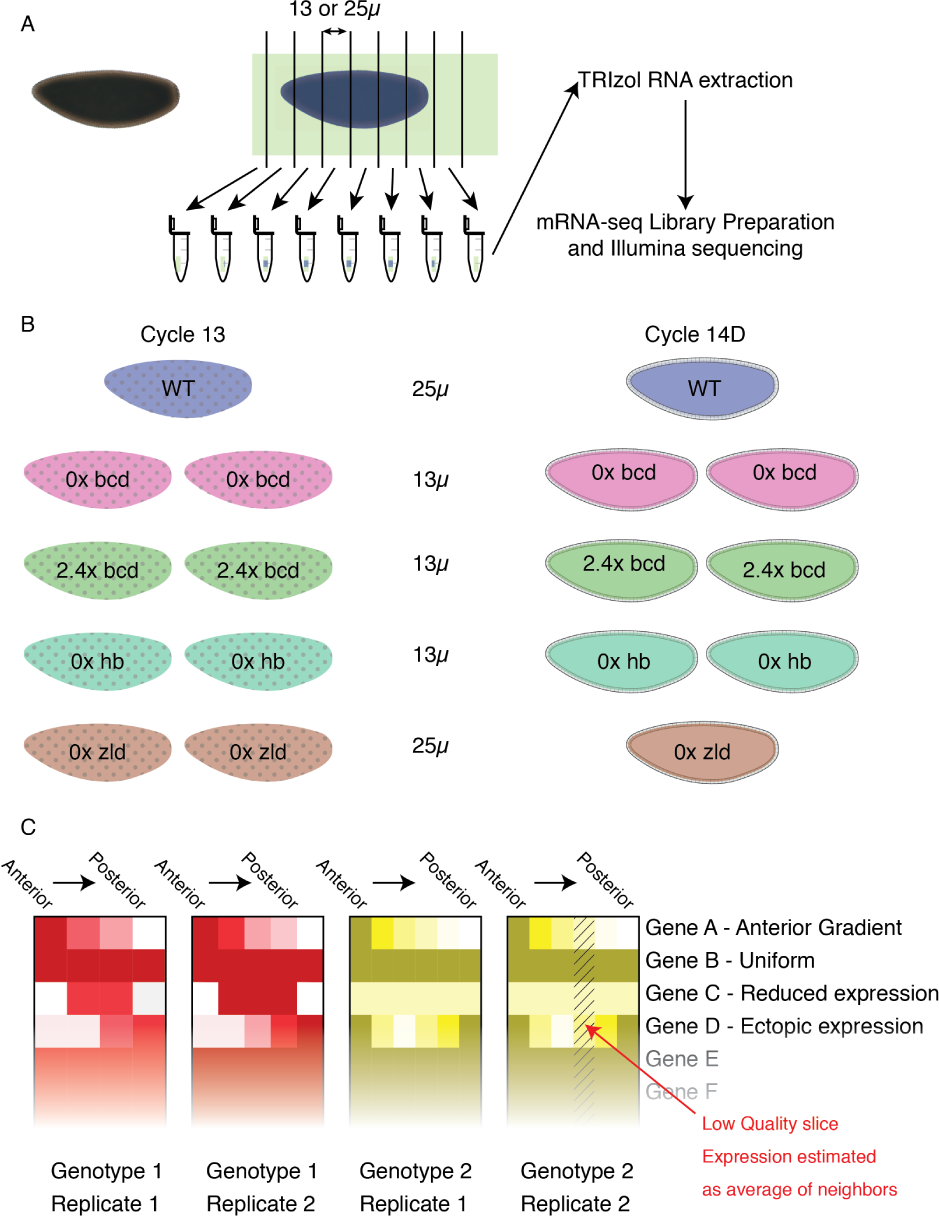
Schematic of experimental approach. A) We fixed embryos in methanol, then selected individual embryos at the correct stage, aligned them in sectioning cups, and sliced to the indicated thickness. We extracted RNA from individual slices, prepared barcoded libraries, then pooled them prior to sequencing. B) Overview of the mutant genotypes used. Two replicates per time point at two time points, based on nuclear density and morphology. C) Cartoon of heatmaps. Each genotype is assigned its own color (matching those in B), with darker colors representing higher expression and white representing no expression detected in that slice. Each boxed column represents a single individual, and within that column, slices are arranged posterior to the left and anterior to the right.

In order to show the range of patterning differences observed, we generated heatmaps of all the gene expression present in the dataset (Fig. 2). Of the 7104 genes with at least 15FPKM in at least one slice, approximately 3000 had uniform expression in all the wild-type embryos that was not greatly perturbed in any of the mutants. The total number of expressed genes is very consistent with previous estimates of the number of maternally deposited and zygotically transcribed genes (Lott et al. 2011).

**Figure 2 (preceding page).**
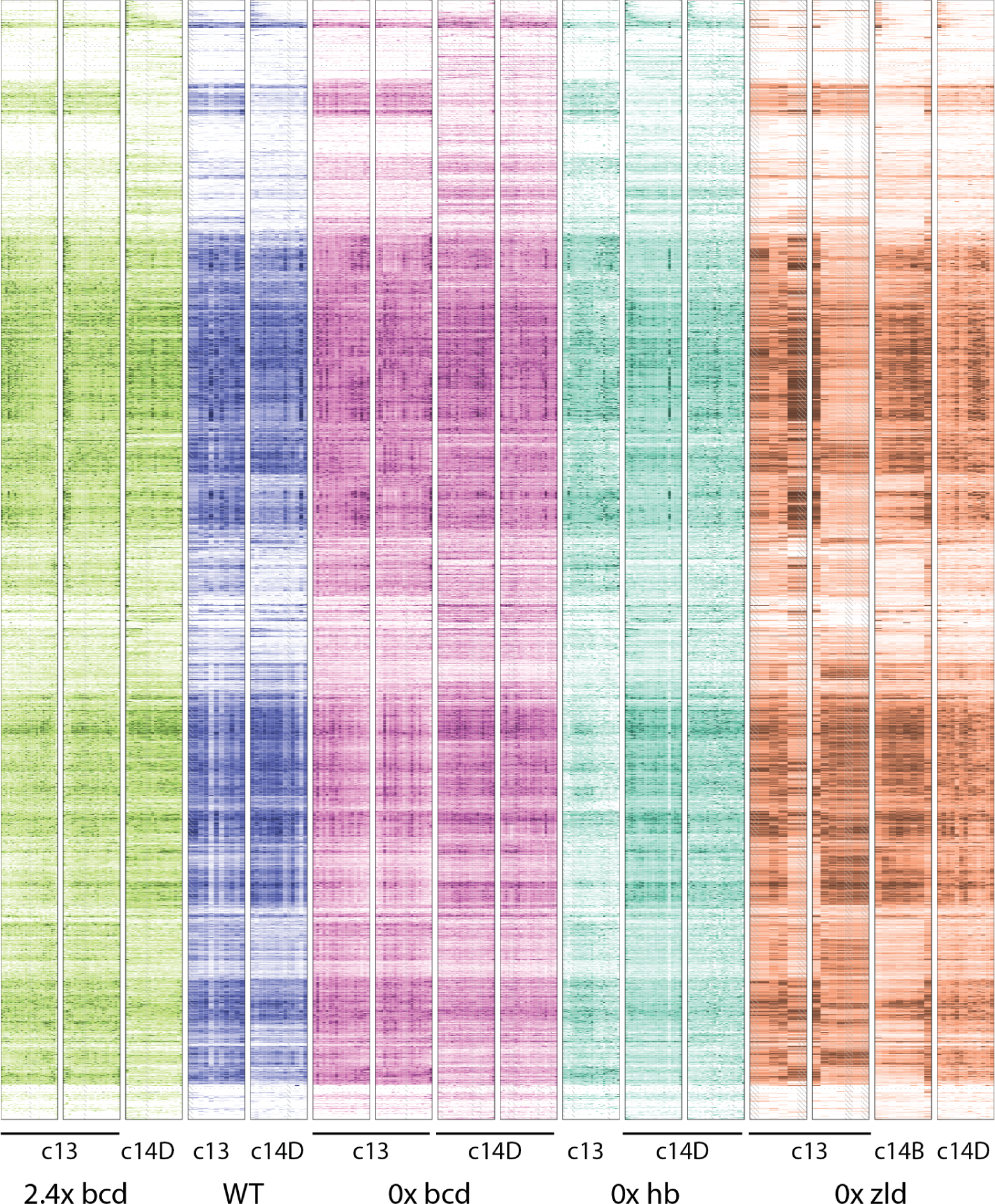
Heatmaps of gene expression patterns for all expressed genes. Each individual embryo is represented by one boxed column in the heatmap. Within a column, slices are arranged anterior to the left, and posterior to the right. Each embryo is colored according to genotype, with green for the bcd over-expression, blue for wild-type, magenta for bcd knockdown, cyan for hb knockdown, and orange for the *zld* mutant. Within a genotype, darker colors correspond to higher expression and white to zero expression, on a linear scale normalized for each gene separately to the highest expression for that gene in the embryo or to 10FPKM, whichever is greater. Slices that did not match quality control standards are replaced by averaging the adjacent slices, and are marked with hash marks. Rows are arranged by using Earth Mover Distance to perform hierarchical clustering, so that genes with similar patterns across all of the embryos are usually close together.

The set of genes with anterior or posterior localization recapitulate the known literature (Staller, Fowlkes, et al. 2015) and general expectations in the *bcd*‐ case: those expressed in the anterior typically lose expression (Fig. 3A), and those in the posterior also frequently gain an expression domain in the anterior (Fig. 3B). Surprisingly, most of these patterns are qualitatively unaffected in the other mutants. In the absence of *zld,* most of these genes are able to retain the proper anterior patterning (although they may have differences in expression levels). Similarly, these genes seem not to be strongly dependent on maternal *hb* for patterning information, with most genes retaining a distinct anterior expression domain. As described in Liang et al. (2008), there are some genes that are normally ubiquitously expressed in the wild-type that become localized to the poles in the zld- embryo (Fig. 3C).

**Figure 3 (preceding page).**
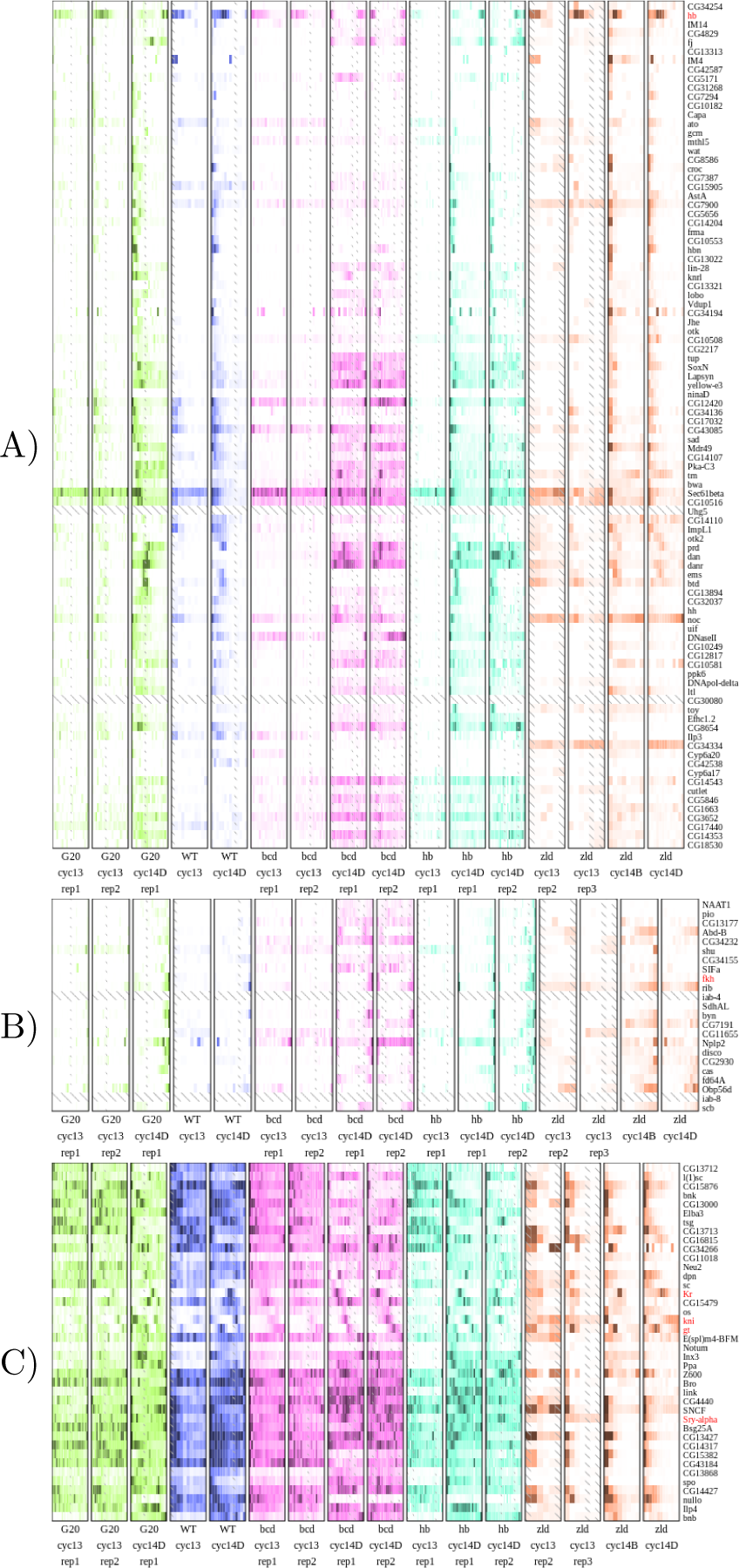
Heatmaps of gene expression patterns for anterior and posterior genes recapitulate expected patterning changes. Each individual embryo is represented by one boxed column in the heatmap. Within a column, slices are arranged anterior to the left, and posterior to the right. Each embryo is colored according to genotype, with green for the *bcd* over-expression, blue for wild-type, magenta for *bcd* knockdown, cyan for *hb* knockdown, and orange for the *zld* mutant. Within a genotype, darker colors correspond to higher expression and white to zero expression, on a linear scale normalized for each gene separately to the highest expression for that gene in the embryo or to 10FPKM, whichever is greater.

### Most spatial patterns are robust to mutation

We next compared expression patterns from each of the mutant lines at late cycle 14 to similar expression patterns in wild-type. Because there are a different number of slices both between the wild type and mutant flies, and between replicates of the mutant flies, we decided to use Earth Mover Distance (EMD) to compare patterns (Rubner, Tomasi, and Guibas 1998). This metric captures intuitive notions about what kinds of patterns are dissimilar, yielding higher distances for dissimilar distributions of RNA, and zero for identical distributions. Patterns were normalized to have the same maximum expression, in order to highlight changes in positioning of patterns, rather than changes in absolute level. In contrast to traditional RNA-seq differential expression metrics, this approach takes advantage of the spatial nature of the data, and with the fine slices, adjacent slices are able to function as “pseudo-replicates”. Adjacent slices are, on average, much more similar than those from farther away in the same embryo (Fig. S1).

The overall level of divergence in pattern across all genes is, in most cases, slightly larger than when comparing nearby time-points in wild-type or replicates of the same genotype and time point (Fig. 4). Notably, the *zld*‐ mutant is more similar to wild-type than the mutants of the other, spatially distributed transcription factors. This suggests that *zld* is a categorically different TF, consistent with its role as a pioneer factor rather than a direct activator. However, the low level of divergence is a reflection of the fact that the majority of genes are not dynamically expressed in either time or space.

**Figure 4.**
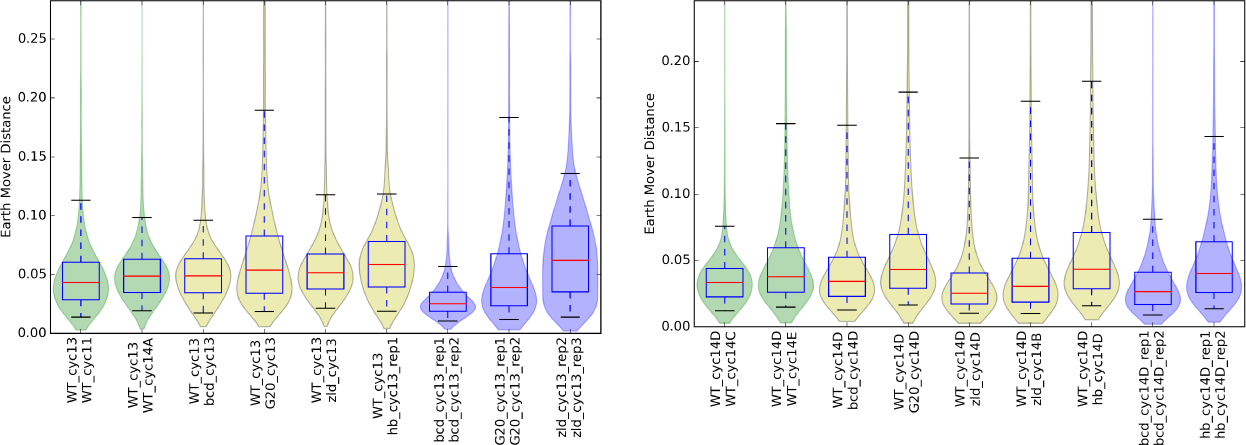
Distributions of patterning differences show that mutants have wide-spread subtle patterning effects and more genes with large patterning differences than replicates. Adjacent time points from the wild-type dataset in Combs and Eisen (2013) are colored green, and replicates of the same genotype and time point are colored blue. Median distances are marked in red.

In order to demonstrate that these mutants are more likely to affect already known BCD regulatory systems, we examined genes that were close to 64 *BCD* dependent enhancers previously identified (H. Chen et al. 2012; Ochoa-Espinosa et al. 2005; Schroeder et al. 2004; Biemar et al. 2005; Hartmann, Reichert, and Walldorf 2001; Kantorovitz et al. 2009; Riddihough and Ish-Horowicz 1991). Although the bulk of these enhancers do not have validated associations with particular genes, we assumed that they would be relatively close to the genes that they drive. Of the 66 genes whose TSS’s were the closest in either direction and within 10kb of the center of the tested CRM, only 32 were expressed at greater than 10FPKM in at least one slice of any of the wild-type embryos. Of these, only 10 had an obvious anterior localization bias (31%), with the majority of the rest being approximately uniformly expressed across the embryo (Fig. 5). The majority of genes with ubiquitious or central localization did not radically change in either the bicoid overexpression or knockdown conditions. As expected, genes with anterior localization suffered a loss of patterning in the depletion mutant, and a posterior shift in the over-expression condition. We assume that genes that are not localized to the anterior are either driven by multiple enhancers, such that loss of expression from one does not severely affect the overall expression, or that they are merely close to the enhancer, but unrelated.

**Figure 5.**
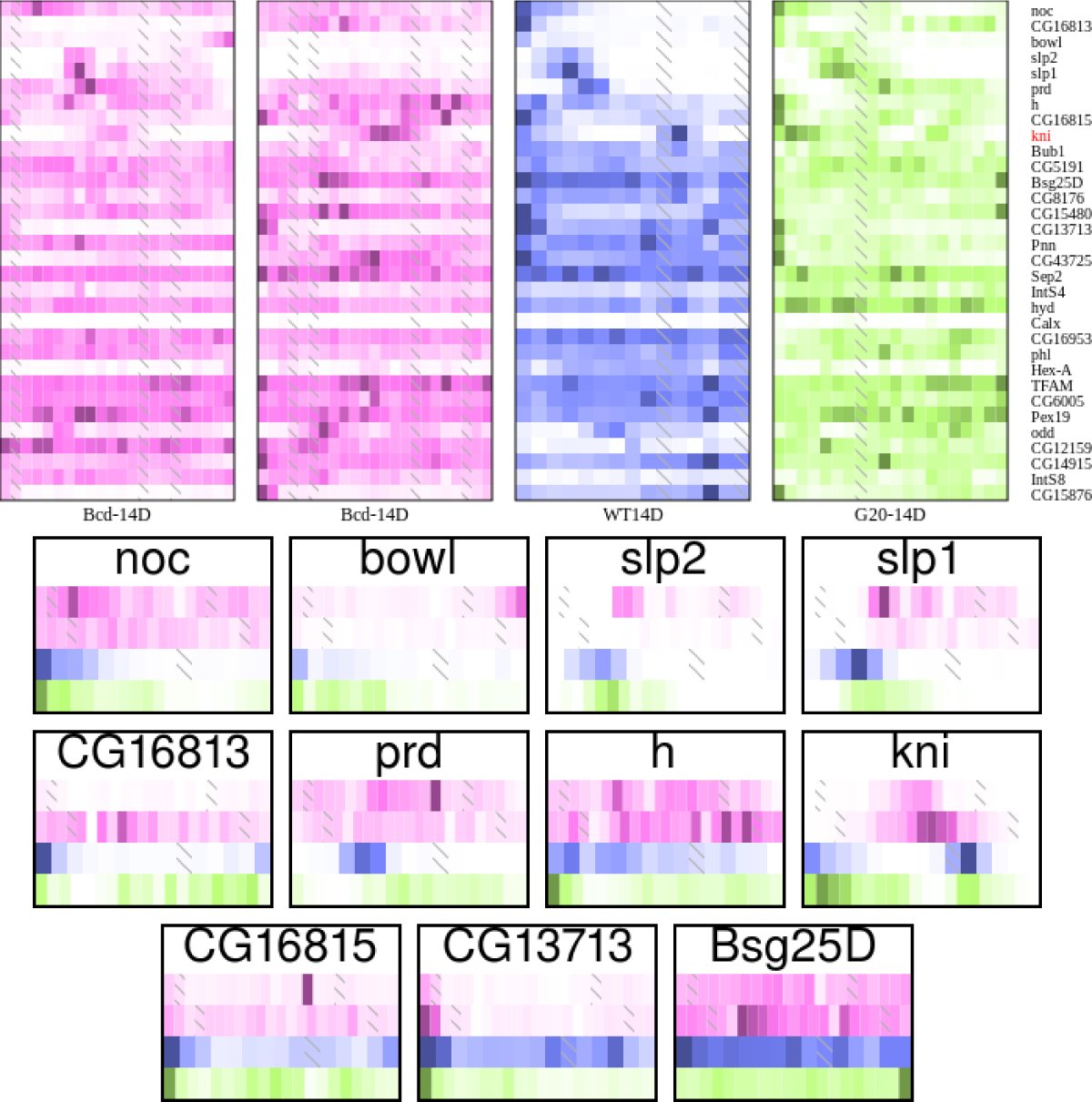
Patterning changes of genes near BCD-dependent enhancers in bcd knockdown and overexpression are clearly visible in anterior-localized genes. A) Each individual is represented in its own heatmap. The magenta heatmaps are from the *bcd*‐ embryos, blue from wild-type, and green from 2.4× *bicoid.* B) Each gene with anterior localized expression in WT, with data from each individual as its own row, to highlight position changes across the mutant genotypes.

**Figure 6.**
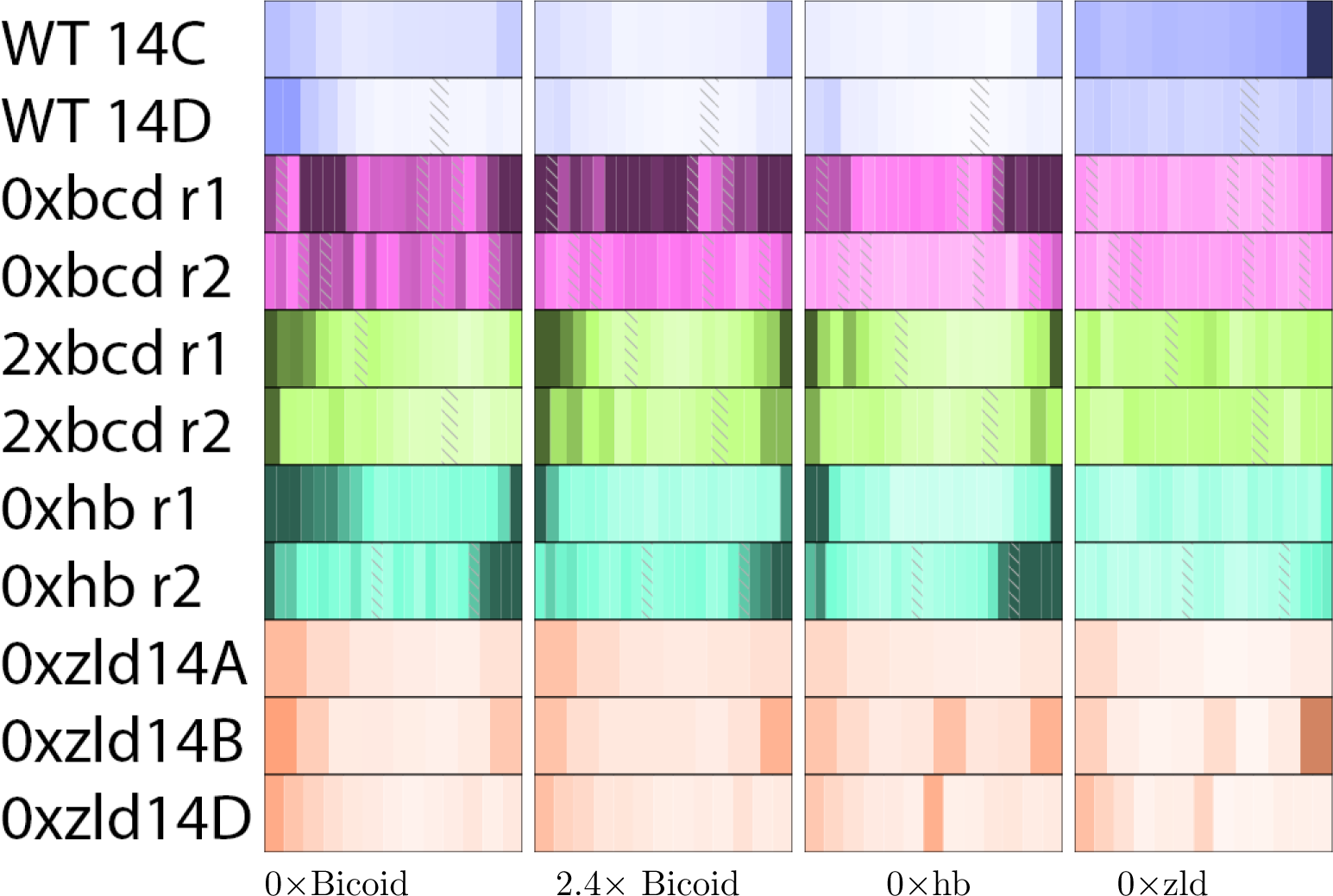
Averaging patterning changes in each genotype recapitulates known gene localization and function. Each individual is represented in its own heatmap. The magenta heatmaps are from the *bcd*‐ embryos, blue from wild-type, and green from 2.4× *bicoid.*

### Effects of TF depletion on patterned genes

We next sought to demonstrate that the technique of cryoslicing mutants is useful for identifying the effects of these early patterning genes. In comparison to fig. 3, where we looked for known patterning changes that we would expect from the literature, we also want to make sure that the largest and most common patterning changes that naturally arise from the data recapitulate the known literature. For each mutant genotype, we

Unsurprisingly, depletion of TFs known to be important for patterning are likely to make an otherwise non-uniform pattern more so (Table 1). Of the 465 genes that have clearly non-uniform patterning in the wild-type at cycle 14D, 12-20% are affected in each depletion mutant, either losing expression entirely or becoming uniform. The over-expression line is at the low end of this range, also at 12%.

**Table 1.** TF depletion is more likely to make a non-uniform pattern uniform than vice versa.

|  | Low Expr to Patterned | Patterned to Uniform | Patterned to Low Expr | Uniform to Patterned |
| --- | --- | --- | --- | --- |
| 2.4×bcd | 343 | 32 | 25 | 64 |
| 0×bcd | 96 | 69 | 40 | 12 |
| 0×hb | 348 | 36 | 20 | 43 |
| 0 ×zld | 71 | 34 | 61 | 28 |

We measured EMD for each gene at cycle 14D in each genotype compared to a uniform distribution. We considered genes uniform if they had an EMD<0.04, and non-uniform if they have an EMD>0.08. We then considered genes with at least 15 FPKM in at least one slice in both wild-type and the mutant line.

However, this is not always simply abrogating expression—a large number of genesseem to have higher expression everywhere. In the case of *bcd* depletion, approximately a third of these cases are genes that are restricted to the anterior in wild-type that become approximately uniform throughout the embryo (Fig. 7). While some of these are due to genes with an early uniform pattern that fails to properly resolve into spatially restricted domains, approximately half are true ectopic expression (Fig. S2).

**Figure 7.**
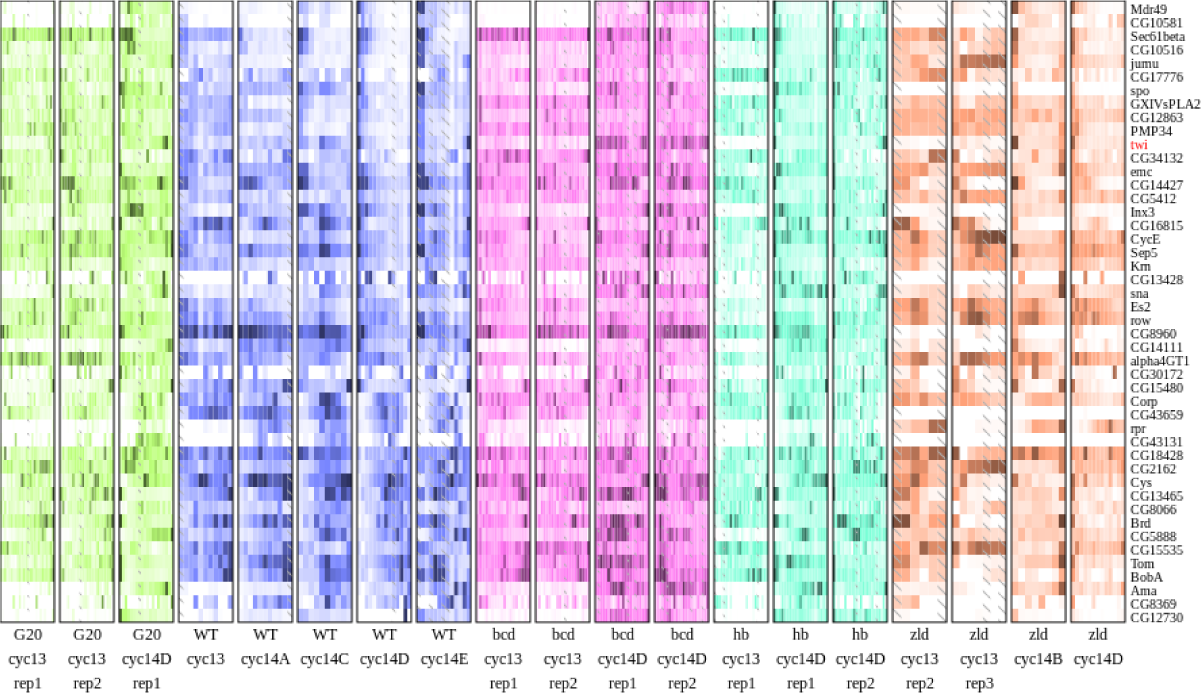
Patterned genes in wild-type that become uniformly expressed are widespread in *bcd*-. Each embryo is normalized independently.

As a first step to identifying likely regulatory motifs, we used binding data for 9 non-pair-rule AP TFs (MacArthur et al. 2009) and for *zld* (Harrison et al. 2011) to search for factors with differential rates of binding among the sets of genes with patterning changes (Table 2). This analysis highlights that *zelda* operates in a qualitatively different manner from the other transcription factors—in it’s absence other TFs are likely to continue expression, though in abnormal patterns. Additionally, *zld* is crucial for maintaining patterned expression, as the most common change is from patterned genes integrating one or more AP factors to minimal overall expression.

**Table 2.**
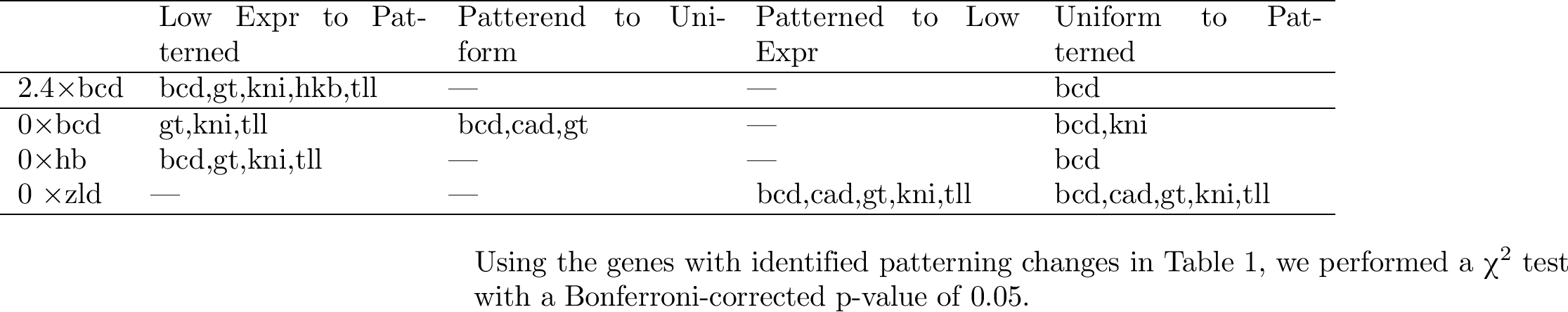
Patterning changes are strongly associated with increased TF binding.

Furthermore, *bicoid* stands out as a major factor involved in AP patterning. In all of the mutant conditions except *bcd* and *zld* depletion, having a *BCD* binding site is associated with an increase in patterned expression. In all of the conditions, a *BCD* binding site is associated with a ubiquitous becoming patterned, and this pattern is often anterior expression.

In addition to patterning changes, some genes with ubiquitous localization actually showed the same response in absolute level as a result of both *bcd* depletion and over-expression. Of these genes, 1002 showed at least 1.5 fold higher expression on both conditions, and 414 showed a 1.5 fold decrease in expression. Such a scenario suggests that these genes are, at wild-type levels, tuned to a particular level of bicoid expression.

It is difficult to reconcile increases of expression in the posterior with any local model of transcription factor action. *BCD* protein is only present at approximately 5nM at 50% embryo length, and negligible levels more posterior (Gregor et al. 2007). It is conceivable that *bcd* activates a repressor gene somewhere in the anterior, which then diffuses more rapidly than *BCD* to cover at least some of the posterior of the embryo. Nevertheless, there have previously been hints that *bicoid* can function as far to the posterior as *hairy* stripe 7 (La Rosee et al. 1997).

### Genes are likely to change in similar ways in different mutant conditions

We next asked whether patterning changes in one genotype could be used to predict whether the pattern changes in another. Therefore, we plotted the EMD between wild-type and the bicoid RNAi line on the X axis, and wild-type to the zelda GLC on the Y axis (Fig. 8). Unsurprisingly, the majority of genes did not change, but of those that did, only a small fraction of them changed in one condition but not the other (the blue and green regions near the axes). We grouped genes according to whether they were in the top 20% of the EMD distribution for each genotype independently, then performed a Pearson's χ^2^ test of independence of change in *bcd*‐ versus *zld-.* The result was highly significant (p<1×10^‐100^), with the largest overrepresentation coming from the case where both changed. Repeating this across all combinations of wild-type and two other mutant genotypes yielded the same results: in every case, there were between 2.2 to 2.7 times as many genes that changed in both categories as would be expected (Fig. S3).

**Figure 8.**
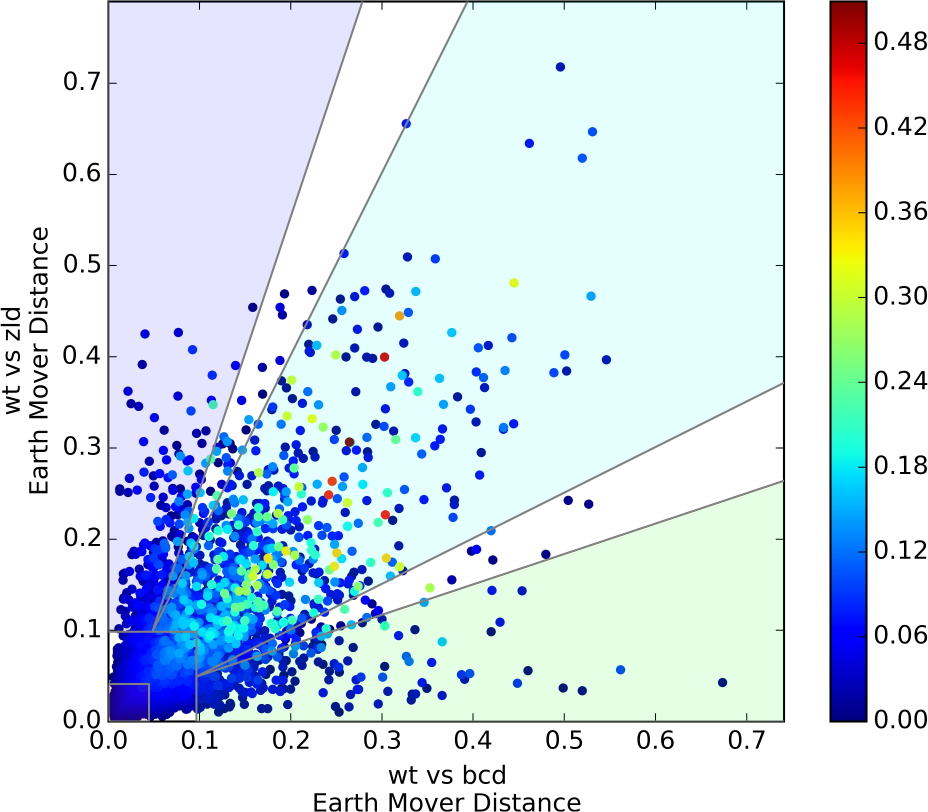
Genes that change in bcdare likely to change in the same way in *zld*-, and vice-versa. Change versus the wild-type is plotted on the x and y axes. Each point is colored according to its ΔD score, calculated in eq. (1), in order to highlight genes that change differently between the two conditions.

Of these genes that do change in both conditions, the majority changed in effectively identical ways. We computed a modified EMD that down-weights genes that are very similar to wild-type in at least one mutant genotype:

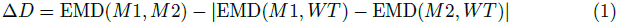

where EMD(x, y) is the Earth Mover Distance between identically staged embryos of genotype x and genotype y. Even among only the set of genes that change in both conditions, ΔD is small (mean of 3.5%, 95th percentile of 11.9%)—equivalent to a shift of the entire pattern by about 1 or 2 slices in either direction. However, there are 13 genes that change differently between wild-type, *bcd-,* and *zld*‐ (ΔD>20%). These genes have noticeably different patterns in all three genotypes (Fig. S4).

### Differential response to mutation is strongly associated with transcription factor binding

We sought to understand what is different about genes with a high ΔD, compared to those that change in response to wild-type, but have a low ΔD (that is, those that change in the same way in response to distinct mutant conditions). We found that genes with a high ΔD score were strongly enriched for a number of TF binding sites (Tables S1 and 3).

**Table 3.** TF binding is enriched near differentially changing genes between WT, bcd-, and zld-.

|  | odds ratio | base freq | p-value |
| --- | --- | --- | --- |
| bcd | 4.69 | 17.40% | 1.2e-09 |
| kni | 4.19 | 3.01% | 0.000176 |
| gt | 3.79 | 19.00% | 2.81e-08 |
| cad | 3.17 | 31.10% | 6.73e-08 |
| kr | 3.1 | 56.58% | 1.6e-07 |
| tll | 2.97 | 7.64% | 0.000597 |
| hb | 2.21 | 39.53% | 0.000143 |
| hkb | 2.1 | 23.86% | 0.00115 |

$\chi^2$ test results for TF binding within 10kb of the TSS for the wild-type/*bcd*-/*zld*- three-way comparison. We examined the top 50 genes by ΔD, compared to the 200 genes closest to the median ΔD of genes that change in response to both mutations. Base frequency indicates the fraction of genes expressed at this time point with at least one ChIP peak for that TF. In this comparison, only Dichaete and zelda binding were not significant at a Bonferroni-corrected p-value of 0.05.

Next, we binned genes by ΔD score, then examined trends in combinatorial transcription factor binding. As ΔD score increases, genes are more likely to be bound by multiple TFs (Fig. 9). Due to the high background rate of binding, assaying the presence of at least 3 factors is not readily able to distinguish between genes with high and low ΔD’s, as nearly 70% of all genes expressed have at least 3 TFs bound. Assaying for the presence of more factors is better able to identify which genes are likely to change, and the top 50 genes all have at least 8 factors bound.

**Figure 9.**
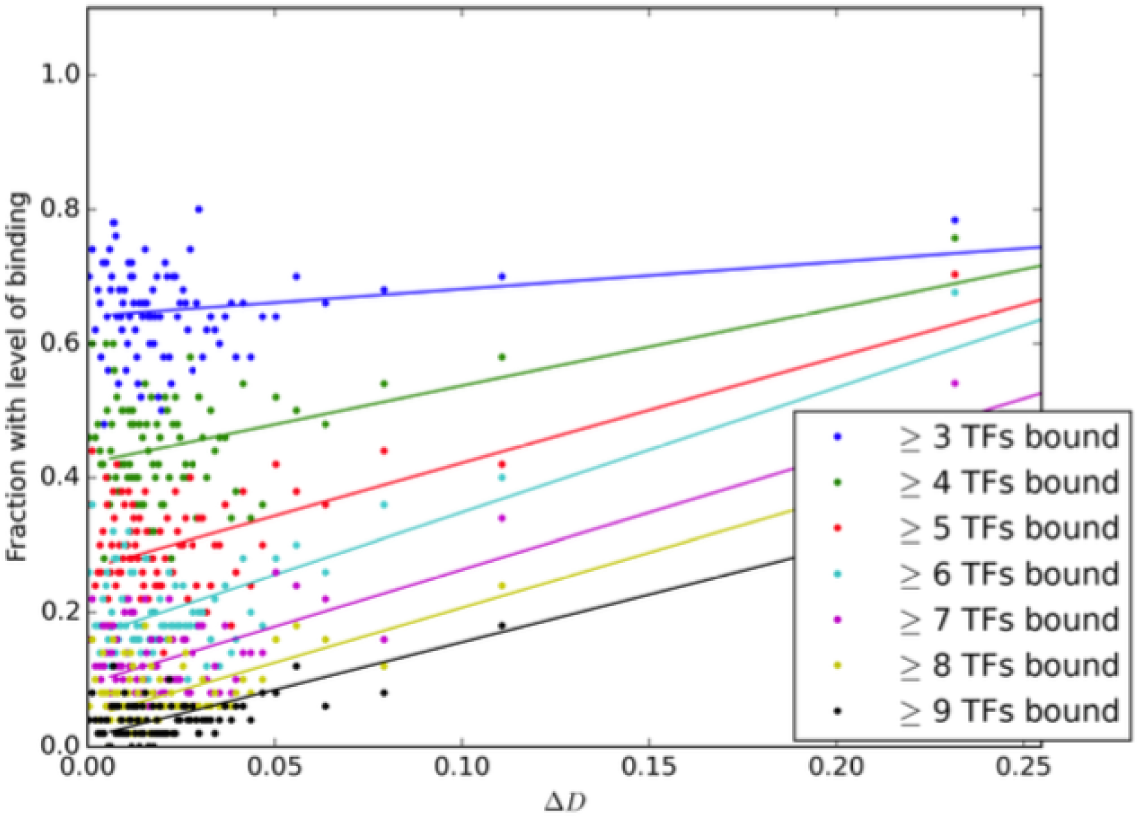
Higher ΔD scores are correlated with increased combinatorial binding. We grouped genes into non-overlapping windows of 50 genes by ΔD score, and calculated the fraction of those genes with at least 3,4, … *etc.* of the 10 early AP TFs bound (including *zld).* We also plotted a simple linear regression on the binned points.

We sought to understand the extent to which genes with the same pattern of upstream regulators had the same responses to perturbation. We grouped genes according to the complement of ChIP-validated TF binding sites near that gene, then examined the patterning changes. Although with 10 different TFs there are potentially over one thousand distinct combinations of binding patterns, in practice the dense, combinatorial patterns found around patterning enhancers reduces this set to a much more manageable 157 different combinations, of which only 52 had at least 30 genes.

Within these sets of genes with similar TF binding profiles, we then asked whether the distribution of patterning changes was, overall, any different from the distribution of patterning changes overall. We performed a KS-test between the distribution of summed EMD scores for the 2.4x *bcd, bcd-,* and *zld*‐ between the set of genes with a given binding pattern, and for all genes overall. We found only 2 binding patterns with a Bonferroni-corrected p-value less than .05. Both of these sets were highly bound, and they were also very similar to each other in their binding, differing only in the presence of a *KNI* site (Fig. 10).

**Figure 10.**
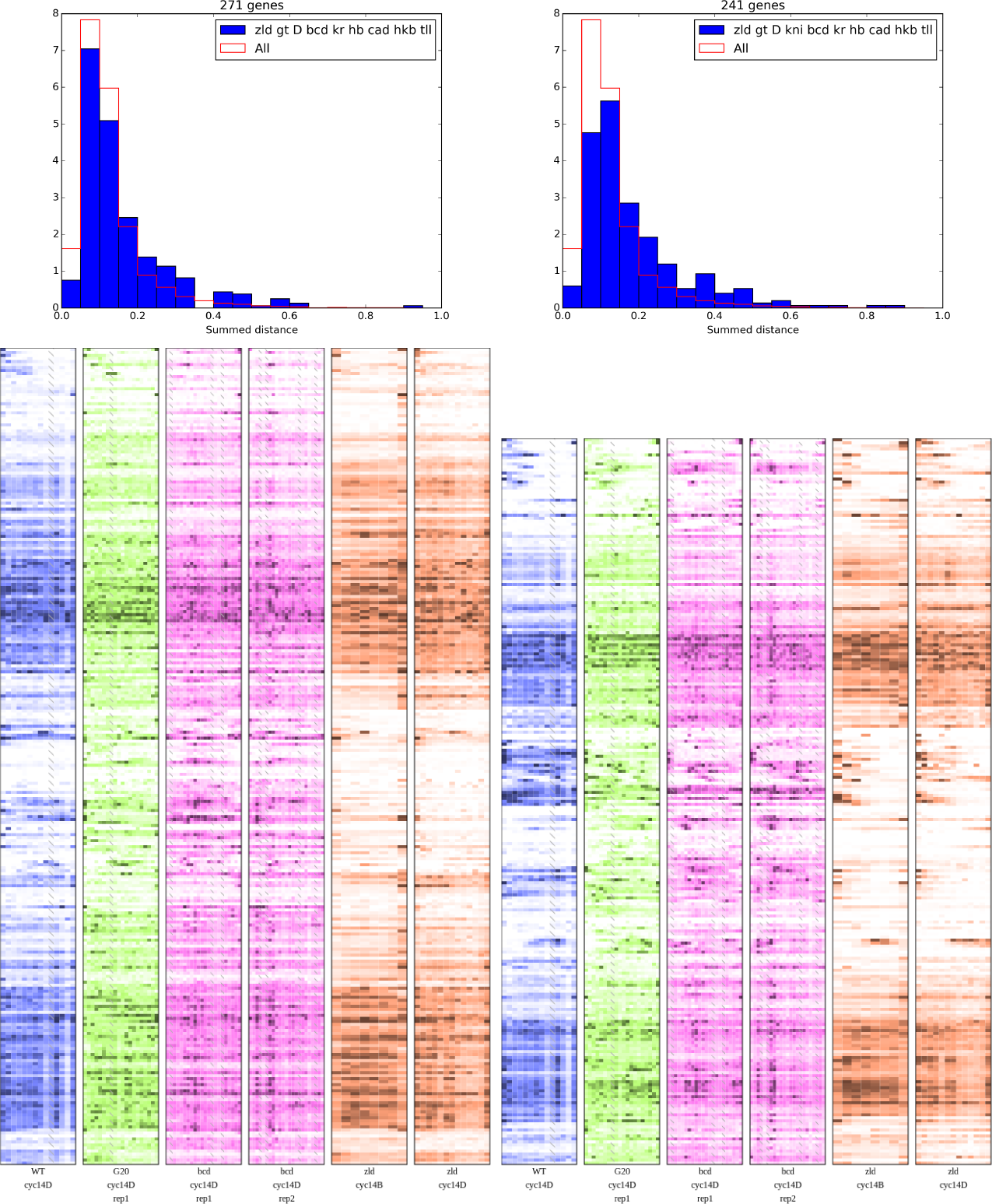
Identical binding patterns have a wide range of patterned responses.

Despite the similar binding patterns near these genes, there is a wide range of responses. The wild-type expression patterns run nearly the complete gamut, including uniform expression, anterior stripes, posterior stripes, and central expression domains. Additionally, the presence of a *KNI* site seems to yield an increased number of genes with an anterior expression domain.

## Discussion

We have generated a dataset that is unparalleled in its coverage assaying patterning changes in mutant conditions. When these patterning mutants have been described previously, either major morphological readouts like cuticle staining or *in situ* hybridization has been used to illustrate the effects on downstream target genes (Driever and Nusslein-Volhard 1988; Liang et al. 2008; Staller, Fowlkes, et al. 2015). However, *in situ* hybridization suffers from a strong selection bias in the genes that are chosen. By assaying spatial differences in the patterning of every gene in the genome, we demonstrate the full effect that these TFs have on developmental gene expression networks.

Despite the importance of the factors we chose for establishing spatially and temporally correct patterning, only a relatively small number of genes have significant expression pattern changes. Many of the targets that do show clearly abnormal expression patterns are, themselves, key transcription factors. This suggests that, even though key, maternally provided patterning factors bind to thousands of places throughout the genome (MacArthur et al. 2009), many of those binding sites are not functional in any meaningful sense. Certainly some of this binding is due to artifacts in the ChIP data, and even reproducible, non-artifactual binding should not be confused with function(Graur et al. 2013; Teytelman et al. 2013). However, the fact that genes near binding sites for multiple factors tend to have more complicated responses to mutation suggests that there is some truth to the idea that gene regulation in complex animals tends to be combinatorial, even if the ChIP data are imperfect.

We were surprised how much proper *bicoid* expression seems to be required for proper patterning at all points along the embryo, not just in the anterior. The fact that *bcd* is normally understood to be an activator, while the plurality of genes with higher, ubiquitous expression in the mutant are normally localized to the anterior in wild-type suggests that this is normally mediated through one or more repressors that depend on *bcd*. As one of three TFs overrepresented at genes with this phenotype, *gt* is likely to be involved in this global derepression, but since it is itself neither ubiquitous throughout the embryo nor universally bound at the genes that change, it is likely not the only player.

The mutants we examined seemed to produce very similar changes in their downstream targets, despite the wild-type TFs having widely varying spatial distributions. Our initial expectation was that there would be many more ways to fail to properly pattern expression, and that different mutations would have different average effects from each other. Indeed, relying on different mutations having different responses has been the key to genetic analysis of fine scale patterns such as the *eve* stripes (Frasch and Levine 1987; Frasch, Warrior, et al. 1988; Small, Blair, and Levine 1996; Andrioli et al. 2002). Although averaging across the most different genes in a mutant genotype does yield different patterns (Fig. 6), for any given gene excursions from the correct spatial expression pattern seems to be largely canalized (Fig. 8). This seemingly-canalized expression change may be a consequence of the types of genes we can easily measure patterning changes among—we cannot resolve individual pair-rule stripes, for instance—so genes with coarser patterns may be more likely to have a single “failure” phenotype, as compared to those with finer patterns, which have more layers of regulation to perturb.

We do recognize a number of distinct limitations of this data set towards predicting gene expression change as a function of mutation. The spatial resolution is still much coarser than *in situ* hybridization based experiments. This is especially concerning near regions where there are fine stripes of expression, which cannot be resolved between adjacent slices, or at regions where there is a transition between expression domains, where it is possible that the slicing axis is not perfectly aligned with the domain border. Finally, it is worth remembering that especially in the later stages examined, the gap gene positions will also be perturbed, so any observed changes in pattern positioning is likely to be a combination of direct effects and downstream effects of the original mutation.

A number of recent studies have used various technical or experimental techniques to improve the resolution of RNA-seq maps of gene expression in developing embryos. Iterated sectioning of different embryos in all three dimensions can be deconvolved to yield estimates of the original pattern (Junker et al. 2014). Similarly, sequencing mRNA from dissociated nuclei allows for the maximum possible spatial resolution, assuming the original location of those nuclei can be estimated (Satija et al. 2015; Achim et al. 2015). While these approaches are worthwhile for establishing a baseline map of expression patterns in wild-type embryos, the expense of sequencing still makes single-dimensional studies worthwhile. Furthermore, the single-cell approaches in Satija et al. (2015) and Achim et al. (2015) require some prior knowledge of spatial gene expression, which may be significantly perturbed in patterning mutants. Other approaches for multiplexed *in situ* profiling of mRNA abundance have been described, but are not yet cheap or reliable enough to be readily useful for screening mutants (Lee et al. 2014; K. H. Chen et al. 2015).

Additionally, the time and expense required for a single individual necessarily means that we have profiled only a small number of individuals. We were therefore careful to choose only highly penetrant mutations for analysis, and to choose individuals at as similar staging as we could. However, even for genes with a consistent, precise time-dependent response between individuals, the differences in staging are likely to be a significant contributor to variation. Furthermore, we only examined two relatively distant time points in this study (approximately 45 minutes apart), making comparisons across time fraught at best.

Nevertheless, this experiment suggests a number of genes for more detailed follow up studies. As our predictive power for relatively well-studied model systems, such as the *eve* stripes improves, it will be especially important to take these insights to other expression patterns in the embryo. The risk of over-fitting increases with the depth of study of any particular model system, even if any given study is relatively well controlled. Therefore, by demonstrating that particular insights hard-won in these model systems are broadly applicable, we can gain some confidence in the results, and we approach having a rigorous, broadly applicable predictive model of gene regulation.

Ultimately, we believe more datasets addressing chromatin state in response to different conditions will be necessary for accurate prediction of spatial responses to mutation. In a ChIP-seq dataset on embryos with different, uniform levels of *bicoid* expression, hundreds of peaks seem to vary with differing affinities to *BCD* protein (Colleen Hannon and Eric Wieschaus, personal communication, March 2015). The zygotically expressed genes near these differential peaks also have different spatial localization in wild-type, and different average responses to the mutants presented here. In addition to spatially resolved expression measurements, spatially resolved binding and chromatin accessibility data will likely be necessary. While ChIP-seq experiments currently require several orders of magnitude more input material than can be reasonably collected from spatially resolved samples, recent methods developments in measuring chromatin accessibility have shown that it is possible to collect data from as few as 500 mammalian nuclei (Buenrostro et al. 2013). A similar amount of DNA is present in a single *Drosophila* embryo, which suggests that spatially resolved chromatin accessibility data may be achievable.

## Materials and Methods

### Fly lines, imaging, and slicing

Zelda germline clone flies (w zld-FRT/FM7a; His2Av RFP) were a gift of Melissa Harrison, and were mated and raised as described previously. Embryos were collected from mothers 3-10 days old.

The construction of the *bcd* and *hb* RNAi flies has been described previously (Staller, Yan, et al. 2013) and were obtained from the DePace Lab at Harvard Medical School. Briefly, we generated F1s from the cross of maternal tubulin Gal4 mothers (line 2318) with UAS-shRNA-bcd or UAS-shRNA-hb fathers (lines GL00407 and GL01321 respectively), then collected embryos from the sibling-mated F1s. In order to take advantage of the slowed oogenesis and resulting greater RNAi efficiency, we aged the F1 mothers for approximately 30 days at 25°C.

The *bcd* overexpression lines were a generous gift of Thomas Gregor at Princeton University. We used line 20, which has 2.4× wild-type levels of eGFP-bcd fusion. Flies were kept in uncrowded conditions, and embryos were collected at 25°C from 3-7 day old mothers.

We washed, dechorionated, and fixed the embryos according to our standard protocol (see (Combs and Eisen 2013)), incubated in 3 μM DAPI for 5 minutes, washed twice with PBS, and then imaged on a Nikon 80i microscope with a Hamamatsu ORCA-Flash4.0 CCD camera. We did not DAPI stain the *zld*‐ embryos because they had a histone RFP marker. After selecting embryos with the appropriate stage according to density of nuclei in histone-RFP or DAPI staining and membrane invagination for the cycle 14 embryos, we washed embryos with methanol saturated with bromophenol blue (Fisher), aligned them in standard cryotome cups (Polysciences Inc), covered them with VWR Clear Frozen Section Compound (VWR, West Chester, PA), and froze them at -80C.

We sliced the embryos as in Combs and Eisen (2013). Single slices were placed directly in non-stick RNase-free tubes (Life Technologies), and kept on dry ice until storage at -80C.

### RNA Extraction, Library Preparation, and Sequencing

We performed RNA extraction in TRIzol as previously (Combs and Eisen 2013). All RNA quality was confirmed using a BioAnalyzer 2100 RNA Pico chip (Agilent).

We generated libraries of the *zld*‐ embryos using the TruSeq mRNA unstranded kit (Illumina). As described previously, we added in 70ng of yeast total RNA as a carrier and performed reactions in half-sized volumes to improve concentration (Combs and Eisen 2013).

We generated libraries from the RNAi and overexpression embryos using the SMARTseq2 protocol; we skipped the cell lysis steps because RNA had already been extracted (Picelli et al. 2014; Combs and Eisen 2015). As described previously, tagmentation steps were performed at 1/5th volume to reduce costs (Combs and Eisen 2015).

### Data analysis and deposition

All data was compared to FlyBase genome version r6.03 (2014_6). Mapping was performed using RNA-STAR v2.3.0.1 (Dobin et al. 2013), and expression estimates were generated using Cufflinks v2.2.1 on only the *D. melanogaster* reads(Trapnell et al. 2012). Reads from Combs and Eisen (2013) were re-mapped to the new genome version. When carrier RNA was used (data from Combs and Eisen (2013) and the *zld*‐ embryos), we discarded as ambiguous reads with 3 or fewer mismatches to prefer one species or the other. The vast majority of mapped reads (>99.99%) of were unambiguous as to the species of origin. After mapping, we removed samples that had fewer than 500,000 *D. melanogaster* reads and samples with less than a 70% mapping rate when no carrier RNA was used; no other filtering or corrections were performed.

Specific analysis code was custom-written in Python. Custom analysis code is available from https://github.com/petercombs/EisenLab-Code. All analyses presented here and all data figures were made using commit 2c144be.

Newly generated sequencing reads have been deposited at the Gene Expression Omnibus under accession GSE71137. Mapped reads, additional files, and a searchable database are available at http://web.stanford.edu/~pcombs/Mutants/.

### In situ hybridization

Probe templates were generated by PCR amplifying cDNA with gene specific T7 promoter fusion primers to cover at least 500 bp of the transcripts. We then made RNA probes using a DIG-labelling kit (Roche), and resuspended in formamide.

Hybridization was performed as described previously (Kosman et al. 2004). All genotypes for each gene were processed in parallel, with hybridization and development times varying by less than 3 minutes.

## Acknowledgments

This work used the Vincent J. Coates Genomics Sequencing Laboratory at UC Berkeley, supported by NIH S10 Instrumentation Grants S10RR029668 and S10RR027303.

---

**Figure S1.**
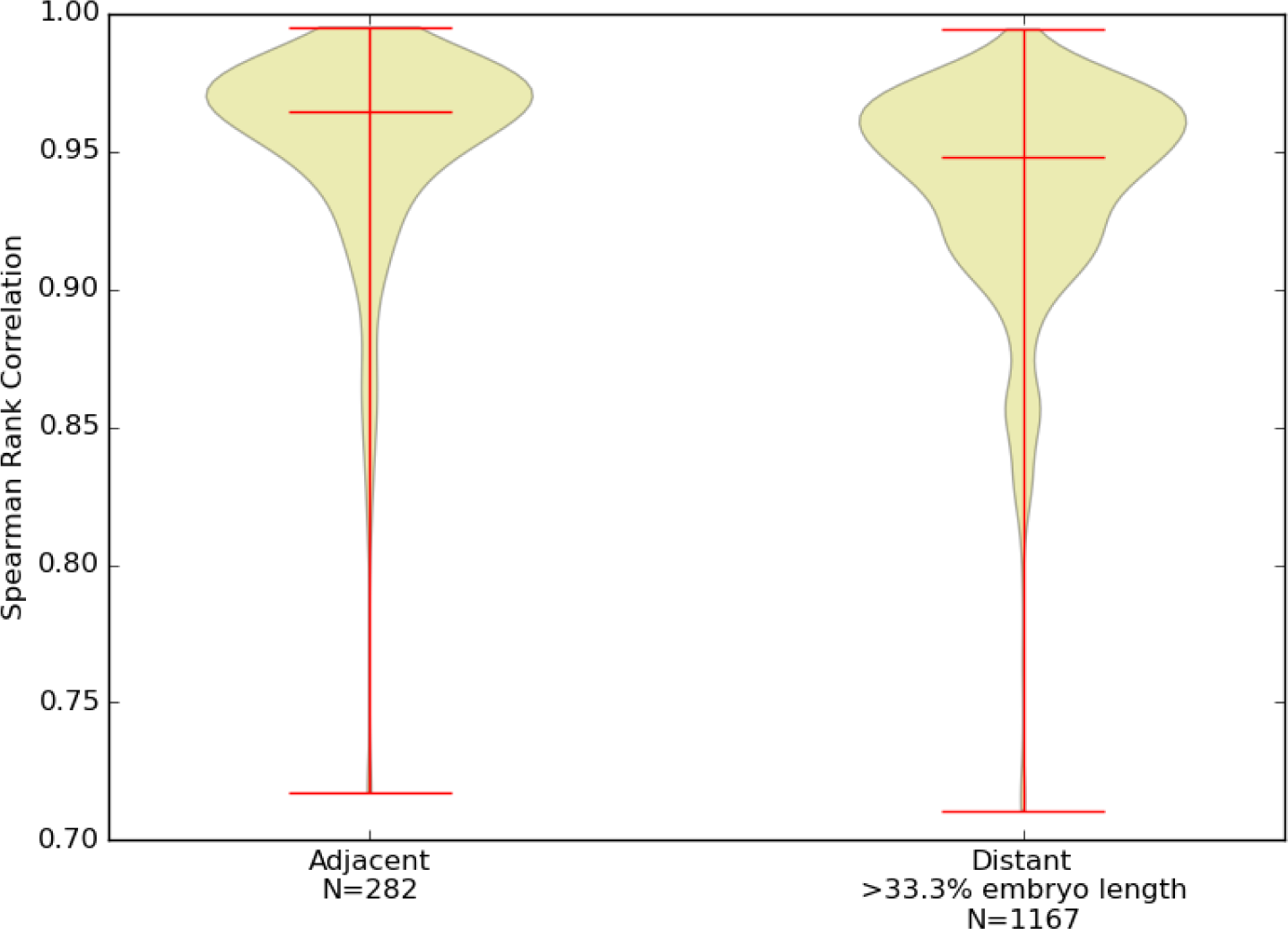
Adjacent slices are more similar than distant ones. Violin plots of the Spearman Rank correlations between adjacent slices and pairs of slices separated by more than one third of the embryo length.

**Figure S2.**
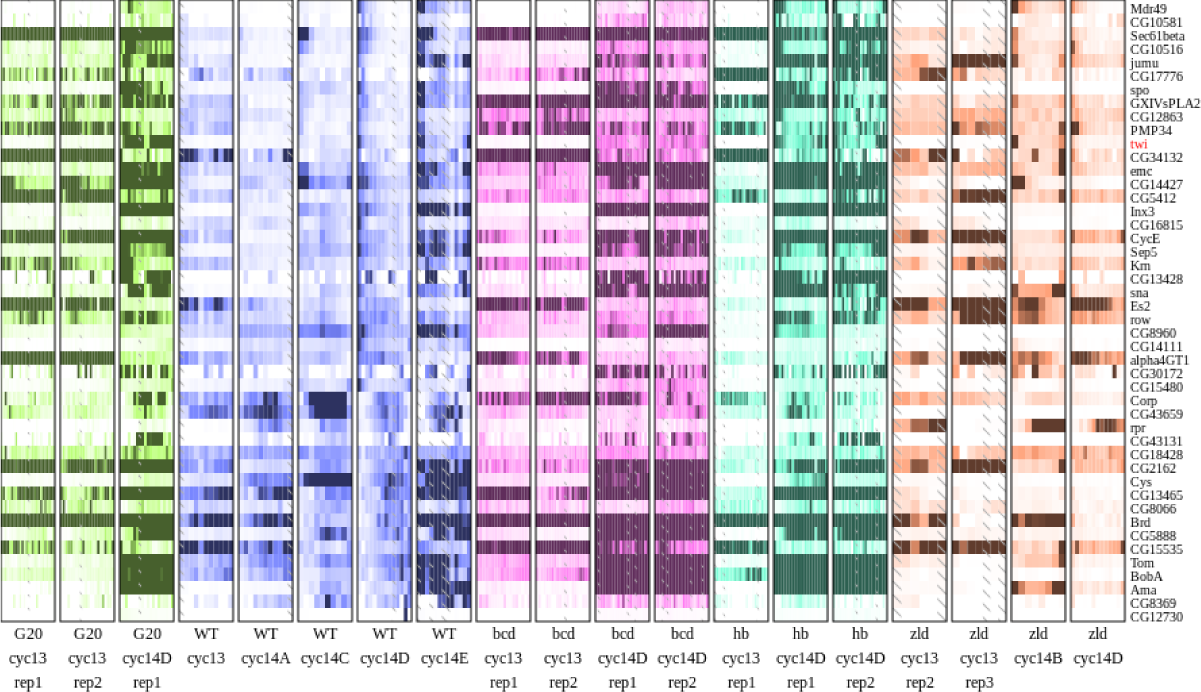
Fig. 7 normalized to expression in wild-type cycle 14D highlight absolute expression level changes. Slices with higher expression are clipped to the maximum expression in wild-type.

**Figure S3.**
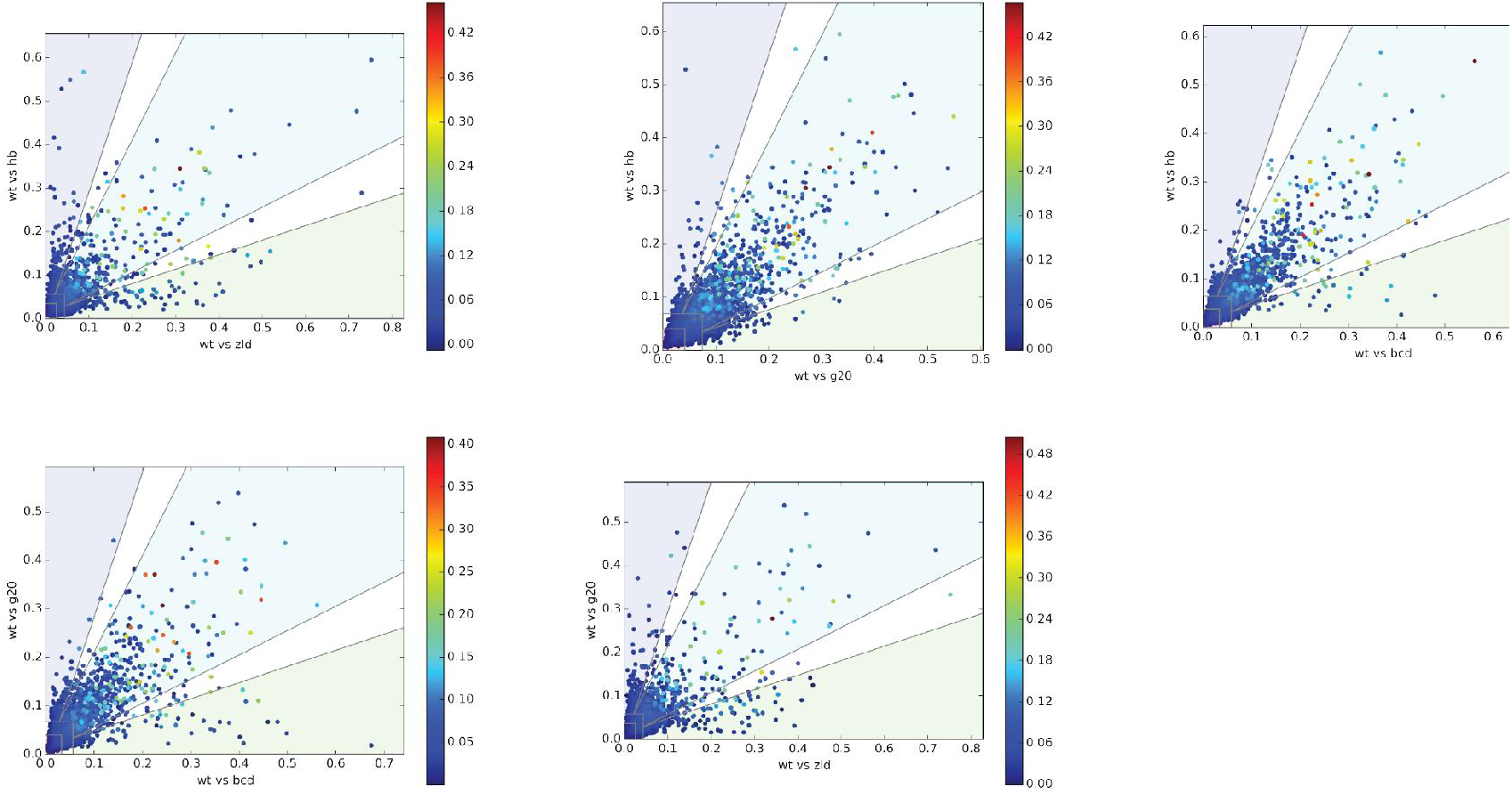
Genes that change tend not to change in only one condition. Three-way comparisons, as in Fig. 8, between wildtype and the remaining combinations of *bcd* depletion, *bcd* overexpression, *hb* depletion, and *zld* depletion.

**Figure S4.**
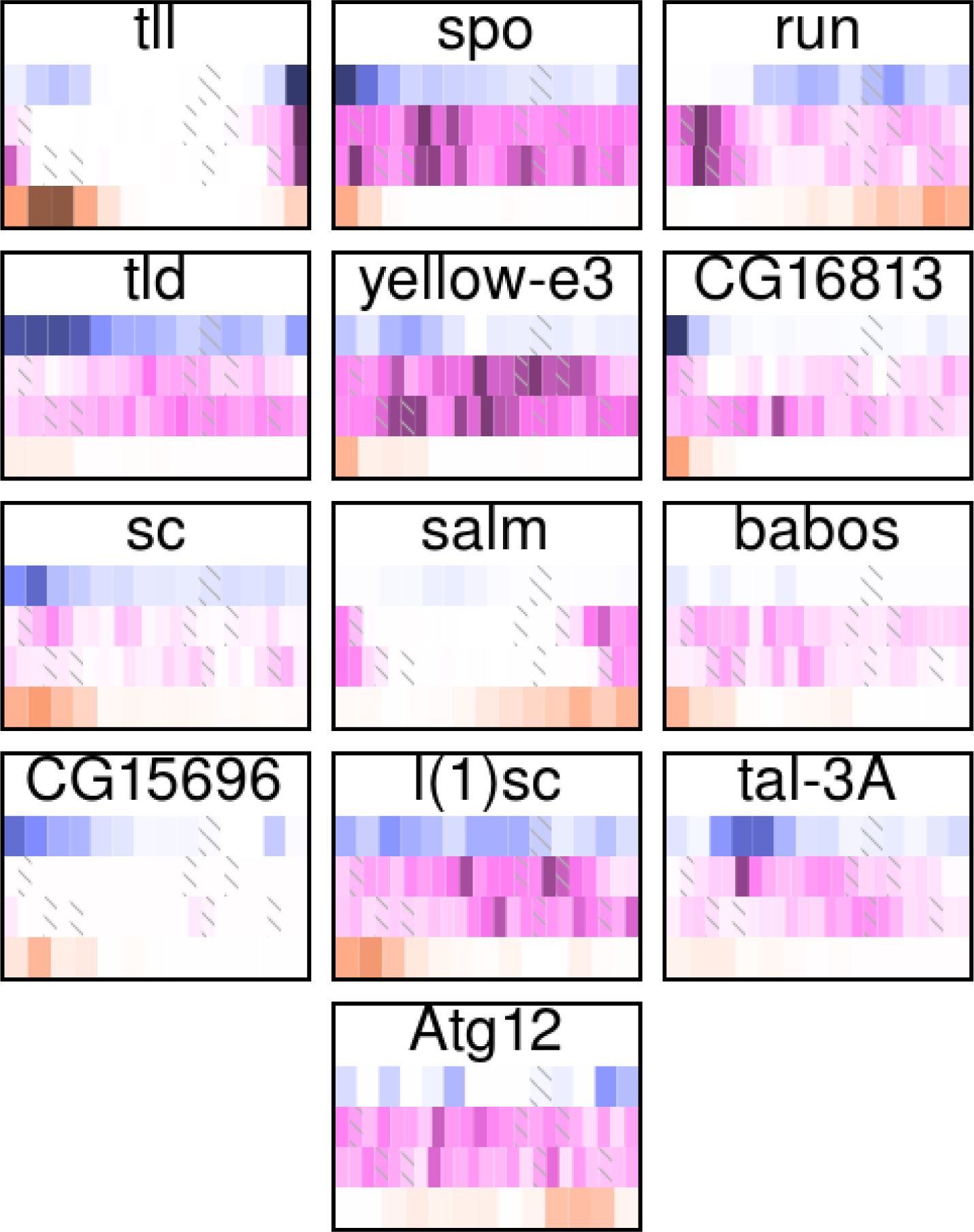
Only a handful of genes change differently between the different conditions. In the WT vs *bcd*‐ vs *zld*‐ three-way comparison, only 13 genes had a ΔD score above 20%. Thumbnails indicate wild-type pattern in blue, *bcd*‐ pattern in both replicates in pink, and *zld*‐ pattern in orange. All expression is scaled to the highest in each individual.

**Table S1.** TF binding is enriched near differentially changing genes across all three-way comparisons.

|  | odds ratio | base freq | p-value |
| --- | --- | --- | --- |
| WT vs *bcd*- vs $2.4\times bcd$ | | | |
| kni | 16.4 | 3.28% | 8.93e-07 |
| tll | 5.5 | 8.08% | 7.72e-05 |
| bcd | 4.1 | 17.58% | 9.24e-05 |
| gt | 3.47 | 19.55% | 0.00053 |
| hkb | 3.1 | 24.15% | 0.0016 |
| cad | 3.09 | 31.98% | 0.00067 |
| WT vs *bcd*- vs *hb*- |  |  |  |
| kni | 11.0 | 3.34% | 0.00297 |
| tll | 5.31 | 8.15% | 0.000671 |
| bcd | 3.29 | 17.67% | 0.00183 |
| D | 0.281 | 90.70% | 0.00251 |
| WT vs *zld*- vs $2.4\times bcd$ | | | |
| kni | 12.3 | 3.05% | 3.63e-07 |
| tll | 11.3 | 7.67% | 3.91e-11 |
| cad | 7.05 | 31.20% | 1.62e-08 |
| bcd | 6.74 | 17.55% | 7.02e-09 |
| gt | 4.56 | 18.99% | 5.95e-06 |
| kr | 4.56 | 56.67% | 8.79e-05 |
| hb | 3.95 | 39.74% | 4.62e-05 |
| WT vs *zld*- vs *hb*- |  |  |  |
| kni | 9.33 | 3.04% | 0.000162 |
| bcd | 7.82 | 17.51% | 6.34e-10 |
| gt | 6.14 | 19.01% | 7.57e-08 |
| tll | 5.05 | 7.67% | 0.000156 |
| cad | 4.77 | 31.01% | 2.05e-06 |
| kr | 3.41 | 56.61% | 0.00102 |
| hb | 2.84 | 39.68% | 0.00186 |
| WT vs $2.4\times bcd$ vs *hb*- | | | |
| tll | 18.5 | 8.16% | 1.19e-07 |
| kni | 8.95 | 3.32% | 0.00169 |
| gt | 4.27 | 19.57% | 3.32e-05 |
| bcd | 3.78 | 17.76% | 0.000173 |
| hb | 2.9 | 40.61% | 0.00147 |
| cad | 2.87 | 31.72% | 0.00152 |

$\chi^2$ test results for TF binding within 10kb of the TSS for the indicated three-way comparison. We examined the top 50 genes by $\Delta D$ , compared to the 200 genes closest to the median $\Delta D$ of genes that change in response to both mutations. Base frequency indicates the fraction of genes with at least one ChIP peak for that TF and that are expressed at this time point in all three conditions.

## References

[1] Kaia Achim et al. “High-throughput spatial mapping of single-cell RNA-seq data to tissue of origin.” In: Nature Biotechnology (Apr. 2015).

[2] Luiz Paulo Moura Andrioli et al. “Anterior repression of a Drosophila stripe enhancer requires three position-specific mechanisms.” In: Development 129.21 (Nov. 2002), pp. 4931–4940.

[3] Frédéric Biemar et al. “Spatial regulation of microRNA gene expression in the Drosophila embryo.” In: Proceedings of the National Academy of Sciences 102.44 (Nov. 2005), pp. 15907–15911.

[4] Jason D Buenrostro et al. “Transposition of native chromatin for fast and sensitive epigenomic profiling of open chromatin, DNA-binding proteins and nucleosome position.” In: Nature Methods 10.12 (Dec. 2013), pp. 1213–1218.

[5] Hongtao Chen et al. “A system of repressor gradients spatially organizes the boundaries of Bicoid-dependent target genes.” In: Cell 149.3 (Apr. 2012), pp. 618–629.

[6] Kok Hao Chen et al. “Spatially resolved, highly multiplexed RNA profiling in single cells.” In: Science (New York, N.Y.) (Apr. 2015).

[7] Peter A Combs and Michael B Eisen. “Low-cost, low-input RNA-seq protocols perform nearly as well as high-input protocols.” In: PeerJ 3 (2015), e869.

[8] Peter A Combs and Michael B Eisen. “Sequencing mRNA from cryo-sliced Drosophila embryos to determine genome-wide spatial patterns of gene expression.” In: PLoS ONE 8.8 (2013), e71820.

[9] Alexander Dobin et al. “STAR: ultrafast universal RNA-seq aligner.” In: Bioinformatics (Oxford, England) 29.1 (Jan. 2013), pp. 15–21.

[10] W Driever and C Nusslein-Volhard. “The bicoid protein determines position in the Drosophila embryo in a concentration-dependent manner.” In: Cell 54.1 (July 1988), pp. 95–104.

[11] M Frasch and M Levine. “Complementary patterns of even-skipped and fushi tarazu expression involve their differential regulation by a common set of segmentation genes in Drosophila.” In: Genes & Development 1.9 (Nov. 1987), pp. 981–995.

[12] M Frasch, R Warrior, et al. “Molecular analysis of even-skipped mutants in Drosophila development”. In: Genes & Development 2.12B (Dec. 1988), pp. 1824–1838.

[13] Dan Graur et al. “On the immortality of television sets: “function” in the human genome according to the evolution-free gospel of ENCODE.” In: Genome biology and evolution 5.3 (2013), pp. 578–590.

[14] Thomas Gregor et al. “Probing the limits to positional information.” In: Cell 130.1 (July 2007), pp. 153–164.

[15] Melissa M Harrison et al. “Zelda binding in the early Drosophila melanogaster embryo marks regions subsequently activated at the maternal-to-zygotic transition.” In: PLoS Genetics 7.10 (Oct. 2011), e1002266.

[16] B Hartmann, H Reichert, and U Walldorf. “Interaction of gap genes in the Drosophila head: tailless regulates expression of empty spiracles in early embryonic patterning and brain development.” In: Mechanisms of Development 109.2 (Dec. 2001), pp. 161–172.

[17] Jan Philipp Junker et al. “Genome-wide RNA Tomography in the zebrafish embryo.” In: Cell 159.3 (Oct. 2014), pp. 662–675.

[18] Miriam R Kantorovitz et al. “Motif-blind, genome-wide discovery of cis-regulatory modules in Drosophila and mouse.” In: Developmental cell 17.4 (Oct. 2009), pp. 568–579.

[19] Dave Kosman et al. “Multiplex detection of RNA expression in Drosophila embryos.” In: Science (New York, N.Y.) 305.5685 (Aug. 2004), p. 846.

[20] A La Rosee et al. “Mechanism and Bicoid-dependent control of hairy stripe 7 expression in the posterior region of the Drosophila embryo.” In: The EMBO journal 16.14 (July 1997), pp. 4403–4411.

[21] J H Lee et al. “Highly Multiplexed Subcellular RNA Sequencing in Situ”. In: Science (New York, N.Y.) 343.6177 (Feb. 2014), pp. 1360–1363.

[22] Hsiao-Lan Liang et al. “The zinc-finger protein Zelda is a key activator of the early zygotic genome in Drosophila.” In: Nature 456.7220 (Nov. 2008), pp. 400–403.

[23] Susan E Lott et al. “Noncanonical compensation of zygotic X transcription in early Drosophila melanogaster development revealed through single-embryo RNA-seq.” In: PLoS Biology 9.2 (2011), e1000590.

[24] Stewart MacArthur et al. “Developmental roles of 21 Drosophila transcription factors are determined by quantitative differences in binding to an overlapping set of thousands of genomic regions.” In: Genome Biology 10.7 (2009), R80.

[25] Amanda Ochoa-Espinosa et al. “The role of binding site cluster strength in Bicoid-dependent patterning in Drosophila.” In: Proceedings of the National Academy of Sciences of the United States of America 102.14 (Apr. 2005), pp. 4960–4965.

[26] Simone Picelli et al. “Full-length RNA-seq from single cells using Smart-seq2.” In: Nature Protocols 9.1 (Jan. 2014), pp. 171–181.

[27] G Riddihough and D Ish-Horowicz. “Individual stripe regulatory elements in the Drosophila hairy promoter respond to maternal, gap, and pair-rule genes.” In: Genes & Development 5.5 (May 1991), pp. 840–854.

[28] Yossi Rubner, Carlo Tomasi, and Leonidas J Guibas. “A metric for distributions with applications to image databases”. In: (1998), pp. 59–66.

[29] Rahul Satija et al. “Spatial reconstruction of single-cell gene expression data.” In: Nature Biotechnology (Apr. 2015).

[30] Mark D Schroeder et al. “Transcriptional control in the segmentation gene network of Drosophila.” In: PLoS Biology 2.9 (Sept. 2004), E271.

[31] S Small, A Blair, and M Levine. “Regulation of two pair-rule stripes by a single enhancer in the Drosophila embryo.” In: Developmental Biology 175.2 (May 1996), pp. 314–324.

[32] Max V Staller, Charless C Fowlkes, et al. “A gene expression atlas of a bicoid-depleted Drosophila embryo reveals early canalization of cell fate.” In: Development 142.3 (Jan. 2015), pp. 587–596.

[33] Max V Staller, Dong Yan, et al. “Depleting gene activities in early Drosophila embryos with the “maternal-Gal4-shRNA” system.” In: Genetics 193.1 (Jan. 2013), pp. 51–61.

[34] Wael Tadros and Howard D Lipshitz. “The maternal-to-zygotic transition: a play in two acts.” In: Development 136.18 (Sept. 2009), pp. 3033–3042.

[35] Leonid Teytelman et al. “Highly expressed loci are vulnerable to misleading ChIP localization of multiple unrelated proteins.” In: Proceedings of the National Academy of Sciences of the United States of America 110.46 (Nov. 2013), pp. 18602–18607.

[36] Cole Trapnell et al. “Differential gene and transcript expression analysis of RNA-seq experiments with TopHat and Cufflinks.” In: Nature Protocols 7.3 (Mar. 2012), pp. 562–578.

